# PMIpred: A physics-informed web server for quantitative Protein-Membrane Interaction prediction

**DOI:** 10.1101/2023.04.10.536211

**Authors:** Niek van Hilten, Nino Verwei, Jeroen Methorst, Carsten Nase, Andrius Bernatavicius, Herre Jelger Risselada

**Affiliations:** Leiden Institute of Chemistry, Leiden University, Einsteinweg 55, Leiden, 2333 CC, The Netherlands; Department of Physics, Technical University Dortmund, Otto-Hahn-Strasse 4, Dortmund, 44227, Germany; Leiden Institute of Advanced Computer Science, Leiden University, Niels Bohrweg 1, Leiden, 2333 CA, The Netherlands; Leiden Academic Centre for Drug Research, Leiden University, Einsteinweg 55, Leiden, 2333 CC, The Netherlands

**Author notes:** These authors contributed equally (Niek van Hilten, Nino Verwei).

## Abstract

**Motivation:** Many membrane peripheral proteins have evolved to transiently interact with the surface of (curved) lipid bilayers. Currently, methods to *quantitatively* predict sensing and binding free energies for protein sequences or structures are lacking, and such tools could greatly benefit the discovery of membrane-interacting motifs, as well as their *de novo* design.

**Results:** Here, we trained a transformer neural network model on molecular dynamics data for *>*50,000 peptides that is able to accurately predict the (relative) membrane-binding free energy for any given amino acid sequence. Using this information, our physics-informed model is able to classify a peptide’s membrane-associative activity as either non-binding, curvature sensing, or membrane binding. Moreover, this method can be applied to detect membrane-interaction regions in a wide variety of proteins, with comparable predictive performance as state-of-the-art data-driven tools like DREAMM, PPM3, and MODA, but with a wider applicability regarding protein diversity, and the added feature to distinguish curvature sensing from general membrane binding.

**Availability:** We made these tools available as a web server, coined Protein-Membrane Interaction predictor (PMIpred), which can be accessed at https://pmipred.fkt.physik.tu-dortmund.de.

## 1 Introduction

Peripheral membrane proteins (PMPs) are a class of proteins that transiently adhere to the surface of the biological membranes that encapsulate and compartmentalise cells. These protein-membrane interactions are crucial for the protein’s subcellular localisation and related function, e.g. in membrane remodelling, transport, or lipid metabolism (Monje-Galvan and Klauda [2016], Whited and Johs [2015]).

The membrane partitioning of PMPs is often regulated by distinct protein regions that have evolved to optimally guide these proteins to their functional locations. One striking example is the amphipathic lipid packing sensing (ALPS) motif that facilitates specific binding to curved membranes and not to flat ones. ALPS was first described as part of the ArfGAP1 protein that regulates COPI coat assembly in a curvature-specific manner (Bigay et al. [2003]). With this original ALPS motif as a template, Drin *et al* defined a set of physicochemical criteria (e.g. prevalence of certain residues, net charge, hydrophobicity, hydrophobic moment) that allowed for a bioinformatic screening of protein databases, which accelerated the discovery of many other curvature-sensing proteins (Drin et al. [2007]).

Despite this breakthrough, there are many membrane-active protein motifs that *do* display similar curvature-related behaviour, but that *do not* fulfill Drin’s criteria. For example, the AH repeats of *α*-synuclein specifically bind to curved anionic membranes (Davidson et al. [1998]), but strongly differ from ALPS in their amino acid (AA) composition (Pranke et al. [2011]). Hence, it is suggested that curvature sensing is governed by a subtle balance between the chemical properties of the hydrophobic and the hydrophilic faces (Drin and Antonny [2010]) that is hard to generalise using simple physicochemical descriptors.

The example of curvature sensing shows that it is not trivial to predict which regions of a given protein structure are responsible for membrane interactions; let alone distinguishing curvature specificity at the same time. Previously developed methods include data-informed classifiers that predict which AAs in a protein structure are membrane-penetrating (e.g. MODA (Kufareva et al. [2014]) and DREAMM (Chatzigoulas and Cournia [2022a,b])) or describe the general protein orientation with respect to a (curved) membrane (e.g. PPM3 (Lomize et al. [2022])). Although these methods perform well for their respective protein targets, they lack the quantitative nature that is required to detect the more subtle interactions involved in curvature sensing. In this paper, we present an alternative method that is able to predict protein-membrane interactions for peptides and protein structures from a bottom-up physics-based perspective. Our approach involves a transformer neural network (NN) model that is trained on a large set of molecular dynamics (MD) data on relative binding free energies for short peptides (24 residues) interacting with model membranes. Since our data were initially gathered during the optimisation of curvature sensing (van Hilten et al. [2023]), our model is additionally able to distinguish curvature- (or, equivalently, lipid packing defect-) sensing from general membrane binding. Moreover, because of its physics-informed character, it is – compared to previous data-driven methods – better able to generalise membrane-interaction features across a wide variety of protein families, including lipid kinases, ALPS-containing curvature sensors, *α*-synuclein, and N-BAR proteins. To enable researchers to access our tool without requiring any programming skills, we incorporated it in our Protein-Membrane Interaction predictor (“PMIpred”) web server, that is available at https://pmipred.fkt.physik.tu-dortmund.de.

## 2 Approach

### 2.1 Theoretical background

In previous work, we developed a free-energy calculation method to efficiently quantify curvature-sensing free energy (ΔΔ*F*) for a peptide within (coarsegrained) MD simulations (van Hilten et al. [2022]). ΔΔ*F* is the difference in affinity for a curved/stretched membrane versus a flat equilibrium membrane. Since binding to the lipid packing defects at curved membrane surfaces negates an energetic penalty, ΔΔ*F* has a negative value for most peptides. The larger the magnitude (ΔΔ*F* ≪ 0), the stronger the sensing propensity.

Because ΔΔ*F* is linearly related to the sequence length *L*, we can extrapolate it to our maximal reference length of 24 AAs, which enables fair comparison between peptides of different lengths:

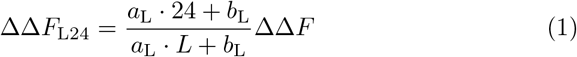

In which *a*_L_ = − 1.03 and *b*_L_ = 3.28. Next, to account for the fact that the majority of biomembranes is negatively charged while the membranes in our simulations were neutral, we can add another correction:

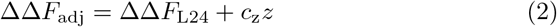

With *c*_z_ = − 0.93 kJ mol^*−*1^, meaning that adding a positive charge *z* reduces ΔΔ*F* by 0.93 kJ mol^*−*1^ and, conversely, adding a negative charge increases it by 0.93 kJ mol^*−*1^.

By combining this approach with an evolutionary algorithm (EA) to efficiently navigate the vast search space (20^24^ possible combinations), we found that the optimal curvature sensor is hydrophobic and bulky, indicating that curvature sensing ΔΔ*F* and general membrane binding Δ*F*_sm_(*R*) – for binding to a vesicle with radius *R* – are strongly correlated:

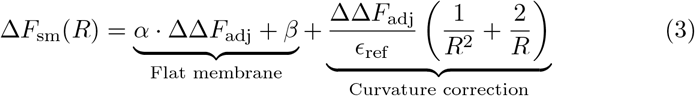

By fitting our data (ΔΔ*F*) to thermodynamic integration simulations that yield Δ*F*_sm_, we found that *α* = 3.83 and *β* = 12.27. The relative strain *ϵ*_ref_ of the target membrane in our MD simulations equals 0.165. The estimation of Δ*F*_sm_ could be alternatively fine-tuned in future generations of the model via incorporation of additional empirically derived algorithmic rules, such as ref. Hristova and White [2005], in order to correct for systematic force field errors. Finally, we theoretically derived that the membrane-binding probability *P*_m_ relates to this Δ*F*_sm_ according to a sigmoid-like switch function:

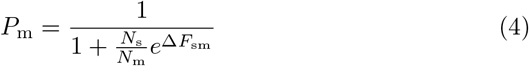

In which the ratio between the number of realisations in solvent (*N*_s_) and on the membrane (*N*_m_) is estimated from typical experimental setups to be 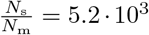. This curve, plotted in Fig. 1A, captures three distinct regimes: non-binders (peptides stay in solution, *P*_m_ → 0), curvature sensors (peptides only bind to curved membranes, *P*_m_ ≈ 0.45), and binders (peptides bind to any membrane, *P*_m_→ 1). Details on the derivations of eq. 1-4 and their constants can be found in our previous publication (van Hilten et al. [2023]).

**Fig. 1.**
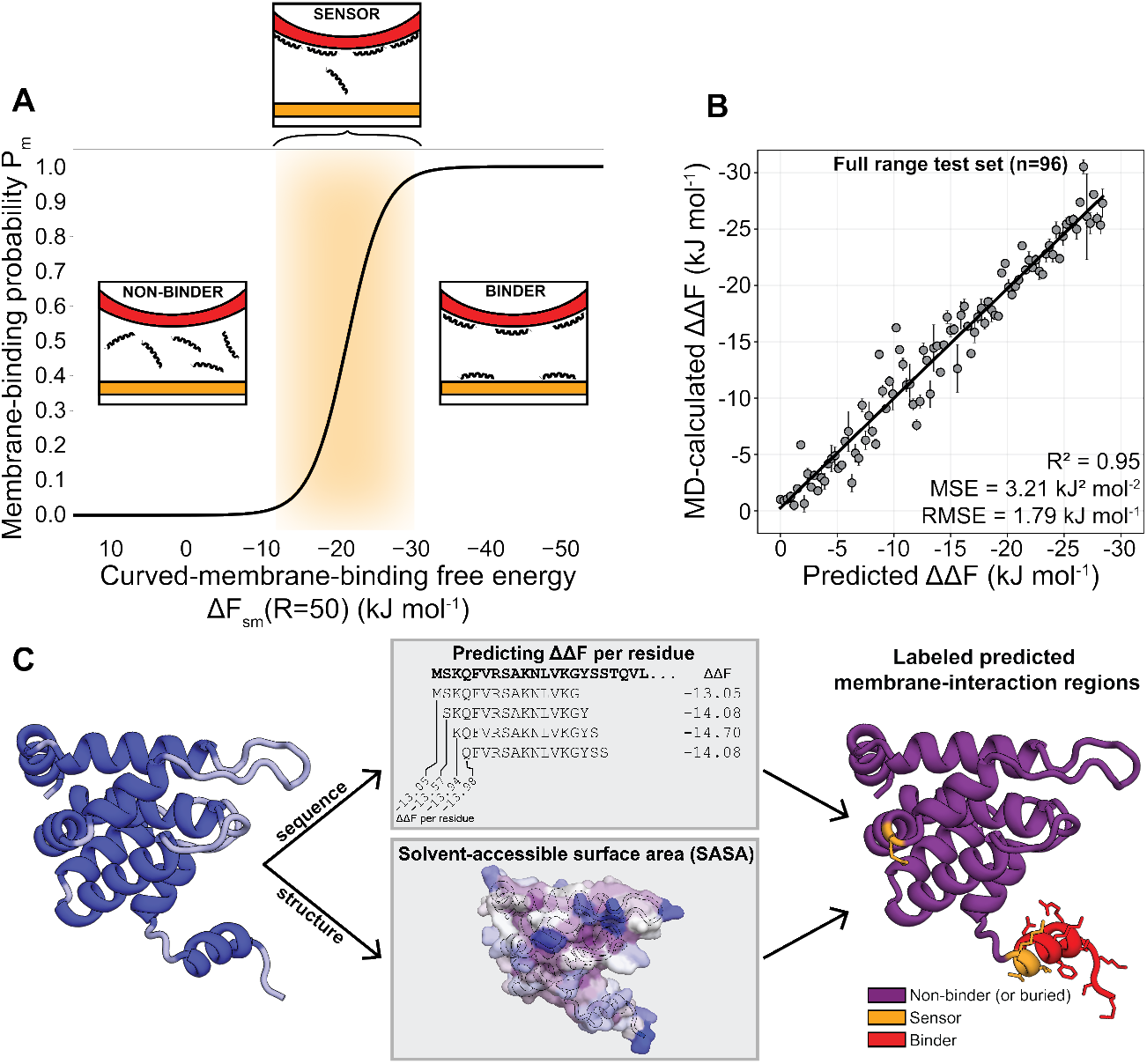
PMIpred’s approach. **A)** The membrane-binding probability *P*_m_ as a function of the membrane-binding free energy Δ*F*_sm_ (at *R* = 50 nm) shows a sharp transition. The orange area marks the ‘sensor regime’ (van Hilten et al. [2023]). Insets show cartoon explanations of the three classes. **B)** Correlation plot for 96 sequences (all 24 residues long, not part of the training data) spanning the full ΔΔ*F* -range. MD-calculated values are averages and standard deviations for 3 independent replicas (500 ns simulation per run, as described in ref. van Hilten et al. [2023]).**C)** Schematic workflow of predicting membrane-interaction regions in protein structures within PMIpred. The trained transformer model predicts ΔΔ*F* (in kJ mol^*−*1^) for overlapping peptide segments from the protein sequence. From the 3D protein structure, per-residue SASA values are calculated. Finally, those two features are combined to yield the final quantitative prediction, as visualised by PMIpred. Non-binding and buried residues are shown in purple. Sensing residues are shown in orange. Binding residues are shown in red. The protein structure shown here is the espin1 ENTH domain (PDB: 5onf).

### 2.2 Transformer model

The randomly initiated evolutionary optimisation of curvature-sensing peptides we performed previously (van Hilten et al. [2023]) yielded a large data set of 53,940 unique peptide sequences (all 24 residues long) and their respective ΔΔ*F* values, as calculated by coarse-grained MD simulations (see S1). An advantage of such a physics-based data set is that it spans the entire applicability domain from ‘inactive’ (ΔΔ*F* = 0.0 kJ mol^*−*1^) to the theoretical optimum (ΔΔ*F*≈− 30.0 kJ mol^*−*1^). Previously, we showed that a convolutional neutral network (CNN) trained on these data is capable of accurately predicting a peptide’s ΔΔ*F* value and classifying its membrane-binding behaviour according to the three aforementioned categories (non-binder, sensor, binder, see Fig. 1A) (van Hilten et al. [2023]). Moreover, it did so across different peptide families and with point-mutation sensitivity.

In the current work, we follow-up on this by employing a transformer model to the same problem. Transformer models use a so-called ‘attention mechanism’ to learn patterns from text input, making them especially useful in natural language processing (Vaswani et al. [2017]). Transformer NNs are able to process whole sentences (or in our case, peptide sequences) at once, which gives them two advantages compared to CNNs and long short-term memory (LSTM) models: (1) they can process input faster, and (2) they are better able to learn long range relations between elements, which is important when – for example – the first and last word of a long input sentence are conceptually and/or grammatically connected.

All details on the transformer architecture, optimisation, and training are described in the supplementary materials, section S1.

Our final transformer model is able to predict the ΔΔ*F* calculated from MD simulations with an expected error (root-mean square deviation) of 1.79 kJ mol^*−*1^, across the full free energy range (Fig. 1B), which is similar to the typical standard deviations we get for the MD simulations in the training data.

### 2.3 Identifying membrane-interaction regions in protein structures

In order to utilise our method to detect membrane-interaction regions in PMPs, we apply a sliding window that divides the full protein sequence into equally sized segments and predicts ΔΔ*F* values for each of them. By default, we consider segments of 15 residues, which is a typical length for a secondary structure element in a protein. A ‘per-residue’ ΔΔ*F* score is obtained by taking the average of the overlapping segment scores at each position (Fig. 1C).

These per-residue values can be interpreted as the contribution of that single amino acid to the overall membrane-binding ability of the protein region it is in. When classifying residues as non-binders, sensors, or binders, we use the ΔΔ*F* -thresholds that follow from the definitions in eq. 1-4: ΔΔ*F >* -6.4 kJ mol^*−*1^ for non-binders; 10.0≤ ΔΔ*F*≤ − 6.4 kJ mol^*−*1^ for sensors, and ΔΔ*F <* -10.0 kJ mol^*−*1^ for binders.

Next, it is important to consider whether a residue is at the surface of the protein structure (e.g. from the PDB or AlphaFold), since this is a prerequisite for it to take part in membrane-binding interactions. Our method accounts for this by calculating the solvent-accessible surface area (SASA) for every individual residue from the protein structure (using the Shrake-Rupley algorithm (Shrake and Rupley [1973]) as implemented in BioPython (Cock et al. [2009]), see Fig. 1C), and then taking a local average of the 9-residue vicinity *n*_*−*4_-*n*_+4_ with the resulting SASA value being assigned to residue *n*.

For intuitive visualisation, the predicted sensing/binding activity can be mapped and colour-coded onto the 3D protein structure (right panel in Fig. 1C). To this end, we use the B-factor field in the output PDB-file that is commonly used for applying colour schemes in molecular visualisation software (e.g. PyMOL (Schrödinger, LLC [2015])). Predicted non-binding (or inaccessible), sensing, and binding regions get assigned B-factors of 0.0, 0.5, and 1.0, respectively.

## 3. Results and Discussion

### 3.1 Predicting membrane interaction for peptides

To apply the trained transformer model on individual peptide sequences, PMIpred allows users to input a string of 7-24 natural AAs (single-letter abbreviations). Also, they need to specify if their target membrane is neutral (e.g. pure POPC) or negatively charged (as is common in nature), see Fig. 2A. In case of the latter, eq. 2 is applied. Upon submission, the web server will produce:

**Fig. 2.**
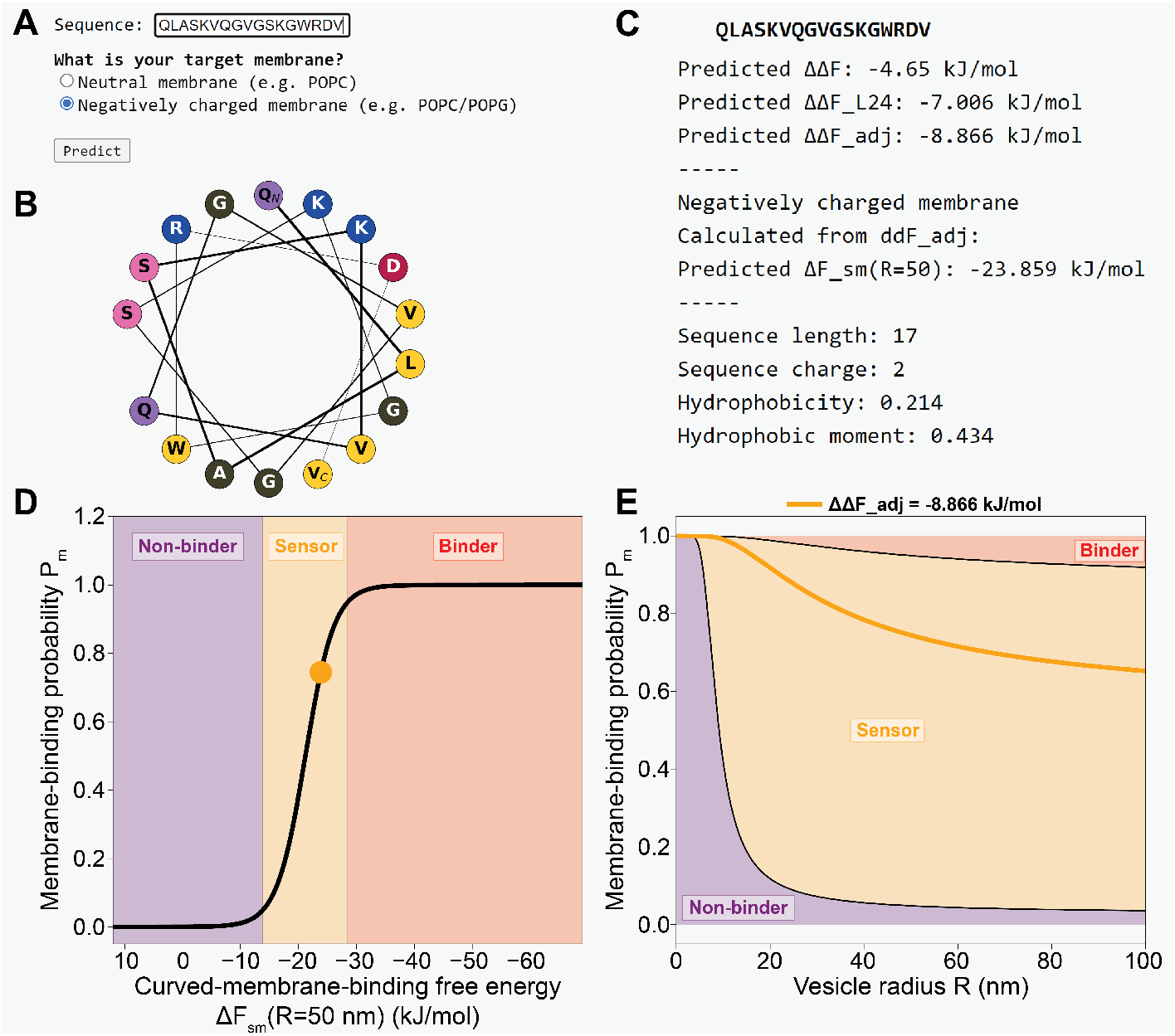
Example output for PMIpred’s peptide module. **A)** Users can input a peptide sequence (part of ArfGAP1’s ALPS2 motif (Mesmin et al. [2007]), in this example) and chose their desired target membrane. **B)** Helical wheel diagram, generated by modlAMP (Müller et al. [2017]). **C)** Predicted free-energy values and physicochemical characteristics. **D)** The position of the input sequence on the non-binding *→* binding continuum. Predicted sensors fall in the orange transition zone. **E)** Membrane-binding probability *P*_m_ as a function of vesicle radius *R*. Again, sensors fall in the orange area.

1. A helical wheel diagram of the input sequence, generated by the modlAMP python module (Müller et al. [2017]) (Fig. 2B). This diagram visualises the spatial orientation of the AA side chains once folded into an *α*-helix. This can be especially useful to, for example, identify a clustering of hydrophobic residues at one side of the helix, which increases the peptide’s amphipathicity and generally favours interactions with lipid bilayer surfaces. Please note that PMIpred – just like the MD simulations that the model is trained on – always assumes an *α*-helical folding for input sequences. Additional MD data on helical and unstructured peptides suggest that membrane-binding free energies are dominated by the general AA composition, and not secondary structure (see S2). Nevertheless, if an input peptide is known to have a non-helical fold (and, potentially, related function), the PMIpred results should be interpreted with caution.

2. Predicted free energies and several commonly used physicochemical descriptors (Fig. 2C). From top to bottom: (1) The NN-predicted curvature-sensing free energy (ΔΔ*F*). The greater the magnitude of |ΔΔ*F* |, the higher the preference for the peptide to bind to (strongly curved) membranes. (2) The length-adjusted ΔΔ*F*_L24_ (eq. 1), that enables fair comparison for peptides with different lengths. (3) The charge-adjusted ΔΔ*F*_adj_ (eq. 2) that additionally corrects for charge-sensing effects on negatively charged membranes. (4) Δ*F*_sm_(*R* = 50), i.e. the predicted membrane-binding free energy to vesicles with a 50 nm radius (eq. 3). Depending on the choice of target membrane (neutral or negatively charged), ΔΔ*F*_L24_ or ΔΔ*F*_adj_ is used for the calculations, respectively. Finally, (5) – and much like tools such as HeliQuest (Gautier et al. [2008]) – PMIpred computes some physicochemical descriptors: sequence length, charge, mean hydrophobicity (using Fauchère & Pliška’s hydrophobicity scale (Fauchère and Pliška [1983])), and Eisenberg’s hydrophobic moment (Eisenberg et al. [1982]).

3.The membrane-binding probability *P*_m_ as a function of the curvedmembrane-binding free energy Δ*F*_sm_ at a 50 nm vesicle radius, as also plotted in Fig. 1A. PMIpred depicts the position of the query sequence on this curve (Fig. 2D).

4. The membrane-binding probability *P*_m_ as a function of vesicle radius *R* (and thus membrane curvature *R*^*−*1^), see Fig. 2E. Peptides that fall in the purple area of this plot are predicted to be non-binders, that prefer to stay in solution regardless of the curvature of available membranes (*P*_m_→ 0). Conversely, the red zone would represent peptides that bind to any membrane, regardless of curvature (*P*_m_→ 1). The orange transition zone is the sensing regime, where the binding probability strongly depends on the membrane curvature. In this plot, a peptide (and its associated ΔΔ*F* value) is represented by a curve, as demonstrated for our example peptide.

### 3.2 Screening proteins for membrane-interaction motifs

As outlined in section 2.3, PMIpred can also be applied to predict which regions in PMPs take part in curvature-sensing or membrane-binding behaviour. To demonstrate this feature, we highlight three example protein structures from different protein families.

#### 3.2.1 ArfGAP1

First, we examined ArfGAP1 (Bigay et al. [2003]), which comprises a structured Zn-finger domain to control GTP hydrolysis in Arf1 on the Golgi membrane (Cukierman et al. [1995]). This catalytic domain is followed by a disordered region that contains two ALPS motifs (Mesmin et al. [2007]), which fold into *α*-helices upon binding to curved membranes. Hereby, they couple the COPI-induced curvature to the catalytic activity and control the transport vesicle budding process. In absence of a PDB entry for the full ArfGAP1 protein, we downloaded the predicted structure from the AlphaFold database (Jumper et al. [2021], Varadi et al. [2022]), which accurately models the distinct structured and unstructured domains (although for the latter, the relative orientation between the loops is likely wrong). We applied our protein screening tool to ArfGAP1 and found that it correctly identifies the two curvature-sensing motifs ALPS1 and ALPS2 (Fig. 3A). Parts of both ALPS motifs were labeled as ‘binders’, although they only slightly surpassed the sensing→ binding threshold at ΔΔ*F* = − 10.0 for most positions (see data repository at https://github.com/nvanhilten/PMIpred). Interestingly, our model also identified a third region (residue 134-151, directly adjacent to the catalytic Zn-finger domain) to potentially contribute to (curvature-specific) membrane interactions. To the best of our knowledge, this segment is not described to be involved in membrane interaction in the current literature, which could mean that either this is a false positive identification, or it is an actual undiscovered membrane-interacting motif that would be interesting to examine further.

**Fig. 3.**
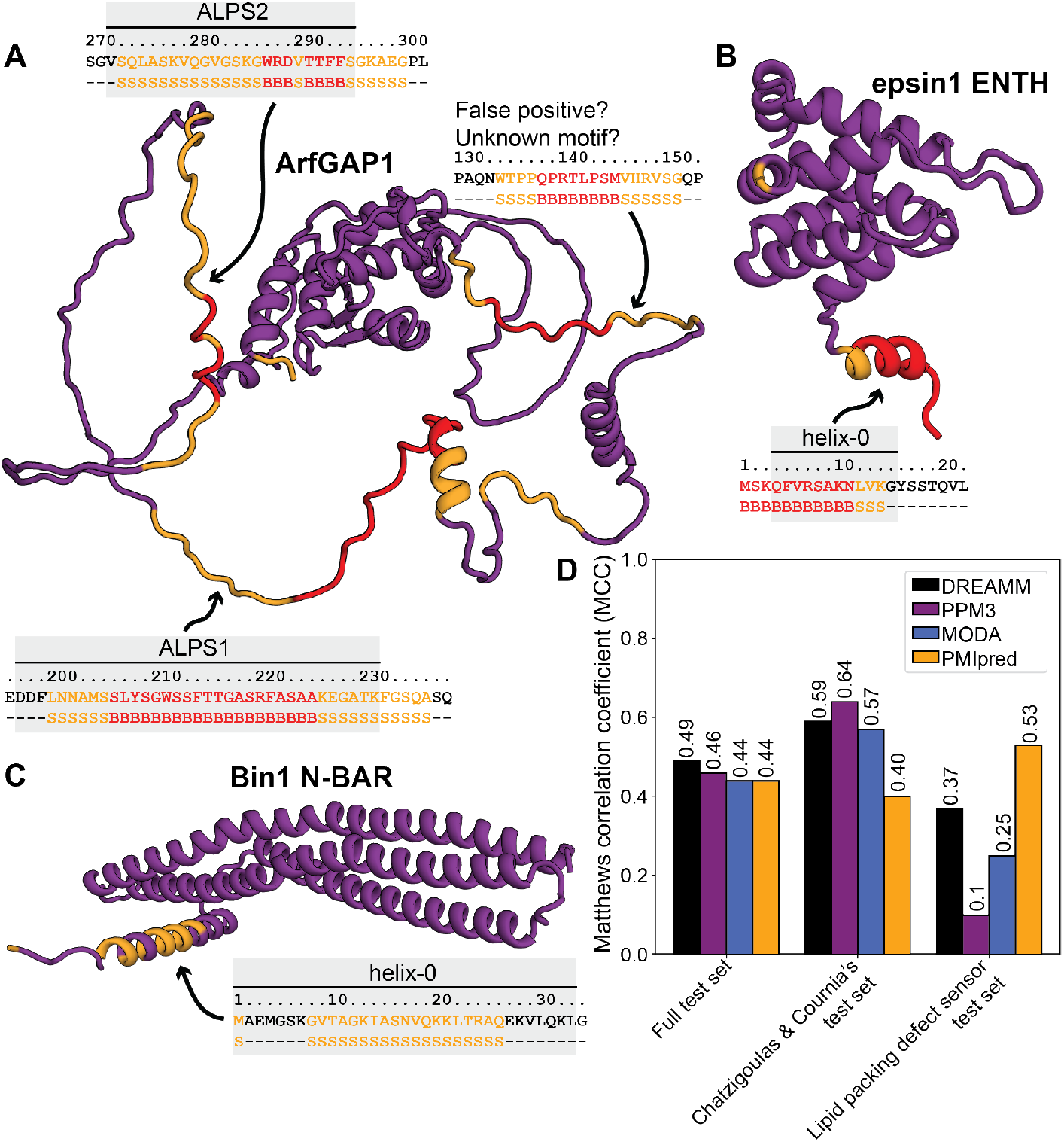
Screening proteins for membrane-interaction motifs. **A-C)** Screening results for ArfGAP1 (**A**, AlphaFold DB: Q8N6T3), the ENTH domain from epsin1 (**B**, PDB: 5onf), and the N-BAR domain from Bin1 (**C**, AlphaFold DB: Q9BTH3). Insets show sequences and corresponding predicted classification of putative active regions. Purple: non-binding or SASA below threshold (-). Orange: curvature sensing (S). Red: membrane binding (B). The known membrane-interaction regions are marked in gray. **D)** Comparing PMIpred to DREAMM, PPM3, and MODA.

#### 3.2.2 Epsin1 ENTH

The second protein we studied is the N-terminal ENTH domain from epsin1, which is responsible for its binding to anionic phosphatidylinositol 4,5bisphosphate (PI(4,5)P_2_) lipids in the inner leaflet of the plasma membrane (Itoh et al. [2001]). Once bound, epsin1 facilitates endocytosis by inducing positive membrane curvature and driving the assembly of clathrin coat components (Ford et al. [2002]). Previous studies have shown that ENTH favours curved membranes (Capraro et al. [2010]), but also interacts with flat membranes to drive curvature by shallow insertion of its N-terminal helix, named helix-0 (Stahelin et al. [2003], Kweon et al. [2006], Lai et al. [2012]). Consistent with these findings, our screening method successfully highlighted helix-0 as a membrane binder (Fig. 3B). Besides the crucial insertion of hydrophobic residues (e.g. L6 (Ford et al. [2002])), helix-0 has a net charge of +4 to complement the negatively charged PI(4,5)P_2_-rich membrane domains that it binds to. The correct identification of this helix in the ENTH domain indicates that, even though the simulations underlying our model were all performed on neutral POPC membranes, the charge correction (eq. 2) we apply is sufficiently accurate to distill these effects.

#### 3.2.3 Bin1 N-BAR

Finally, we screened the N-BAR domain from Bin1, which is a crescent-shaped protein that polymerises to drive the remodelling of membranes into tubes. Hereto, we took the predicted structure of the full Bin1 protein from the AlphaFold database, and isolated the N-BAR domain (residue 1-239). Our screen highlighted part of the N-terminal helix (just like for ENTH, named helix-0), classifying it as a curvature sensor (Fig. 3C). When comparing this to the extensive literature on Bin1’s helix-0, we see that – contrary to our method’s classification – it behaves as a binder in liposome experiments, which show that its membrane binding is not curvature-dependent (Fernandes et al. [2008]) and that both the full Bin1 N-BAR domain and the amphipathic helix0 alone can induce tubulation (Löw et al. [2008]). Given that the BAR domain by itself (so, without helix-0), also induces liposome tubulation (Peter et al. [2004]), the current understanding on this is that helix-0 is responsible for the initial docking and positive curvature induction, which then triggers binding of the full crescent-shaped BAR domain and the subsequent polymerisation of the full protein scaffold (Adam et al. [2015], Simunovic et al. [2015]). Additionally, our screening tool predicted three other sensing/binding domains that were not sufficiently solvent exposed (SASA *<* 0.8 nm^2^) to be highlighted in the results (see data repository at https://github.com/nvanhilten/PMIpred). Two of these domains are indeed located at the membrane-binding face of the concave N-BAR domain.

When taking together our screening results for ArfGAP1, epsin1, and Bin1, we conclude that our method was indeed able to correctly pick out the important regions from these proteins (Fig. 3A-C). However, the sensing/binding classification is not perfect. This likely stems from a combination of inaccuracies in the model and inconsistencies between experimental setups. After all, curvature sensing and membrane binding will be strongly affected by liposome formulations, peptide concentrations, and potential synergistic effects between proteins that may differ between experiments and are not accounted for in our simulations.

#### 3.2.4 Comparison with other tools

Current tools that predict membrane-interaction sites in proteins (DREAMM (Chatzigoulas and Cournia [2022a,b]), PPM3 (Lomize et al. [2022]), and MODA (Kufareva et al. [2014])) produce binary output on the single-residue level (yes or no), whereas PMIpred quantifies membrane-binding *regions*; averaging the free energy over the local surrounding of a given amino acid. This complicates a one-to-one comparison with these qualitative prediction methods on the single-residue level (e.g. as done in ref. (Chatzigoulas and Cournia [2022a])). Therefore, we proceeded to benchmark PMIpred against the other web servers in the following manner: we composed a test set of 27 PMPs, including all 19 proteins from Chatzigoulas & Cournia’s test set (Chatzigoulas and Cournia [2022a]) (mainly comprising lipid kinases and lipid transfer proteins) and 8 known lipid packing defect sensing proteins; see Table S2 for details. Then, we segmented these proteins into consecutive fragments of 15 AAs (matching PMIpred’s default window size), which were marked *positive* if they contained *>* 1 residue that was predicted to be membrane-interacting (either sensing or binding for PMIpred), and *negative* if none of the 15 residues in the fragment was labeled as a ‘hit’. We argue that, although it provides fundamentally different insights than e.g. DREAMM, this ‘protein-region’-way of looking at membrane-binding interactions is more akin to the way a structural biologist may look at a protein structure, with loops or helices (rather than independent individual residues) dynamically inserting into the membrane leaflet.

We performed this segment-based binary classification analysis and calculated the Matthews correlation coefficient (MCC, see S4) for the four methods on the different test sets (Fig. 3D). We observed that all tools score similarly on the full test set (MCC ≈0.5). When only considering the proteins in Chatzigoulas & Cournia’s test set, we see that PMIpred is outperformed, mainly by DREAMM and PPM3, due to the fact that these tools are – by design – focused on predicting the single-residue protein-membrane interactions that are typical for the types of PMPs in this test set. In contrast, PMIpred performed much better when predicting the (often less strictly defined) membrane-interacting regions of lipid packing defect sensors. Details on all results described in this section are provided in Table S3 and the data are accessible at https://github.com/nvanhilten/PMIpred.

In general, we note that the majority of erroneous predictions are false positives rather than false negatives, with a false discovery rate (FDR) of 60% and a false omission rate (FOR) of only 4.4% for PMIpred (and similar values for the other web servers, see S4).

Taken together, we argue that PMIpred’s predictions are generally on par with state-of-the-art tools like DREAMM, PPM3, and MODA. However, we should emphasise that PMIpred additionally provides *quantitative* information on membrane-interaction propensity per residue in actual thermodynamic terms (free energies), whereas DREAMM and PPM3 act as binary classifiers and MODA only yields membrane-interaction ‘likelihoods’, that are hard to interpret biophysically. Moreover, because PMIpred is trained on bottom-up physics-based data, it is likely more self-consistent at generalising those concepts to undiscovered or *de novo* designed protein structures that are highly distinct from typical known PMPs, compared to data-informed models that are trained on protein databases. To this end, we suggest that the physics-informed transformer neural network we developed here may be able to additionally boost the performance of prediction methods based on ensemble machine learning approaches such as DREAMM by serving as an additional, independent (strong) classifier, especially since these approaches are notably orthogonal. Finally, we emphasise that users now have multiple different tools to their disposal and encourage them to use these various approaches to compare results and draw conclusions based on the ensemble of results.

#### 3.2.5 Try it yourself

Our protein screening tool is accessible within the PMIpred web server. Here, users can submit a PDB-file or PDB ID and also tune the size of the sliding window and the SASA threshold value (Fig. 4A). Similar to PMIpred’s peptide module, users can specify if their target membrane is neutral (e.g. POPC) or negatively charged, as is common in biology. Again, if the latter is chosen, charge correction (eq. 2) is applied to the ΔΔ*F* value to account for additional coulomb interactions and concomitant ‘charge sensing’.

**Fig. 4.**
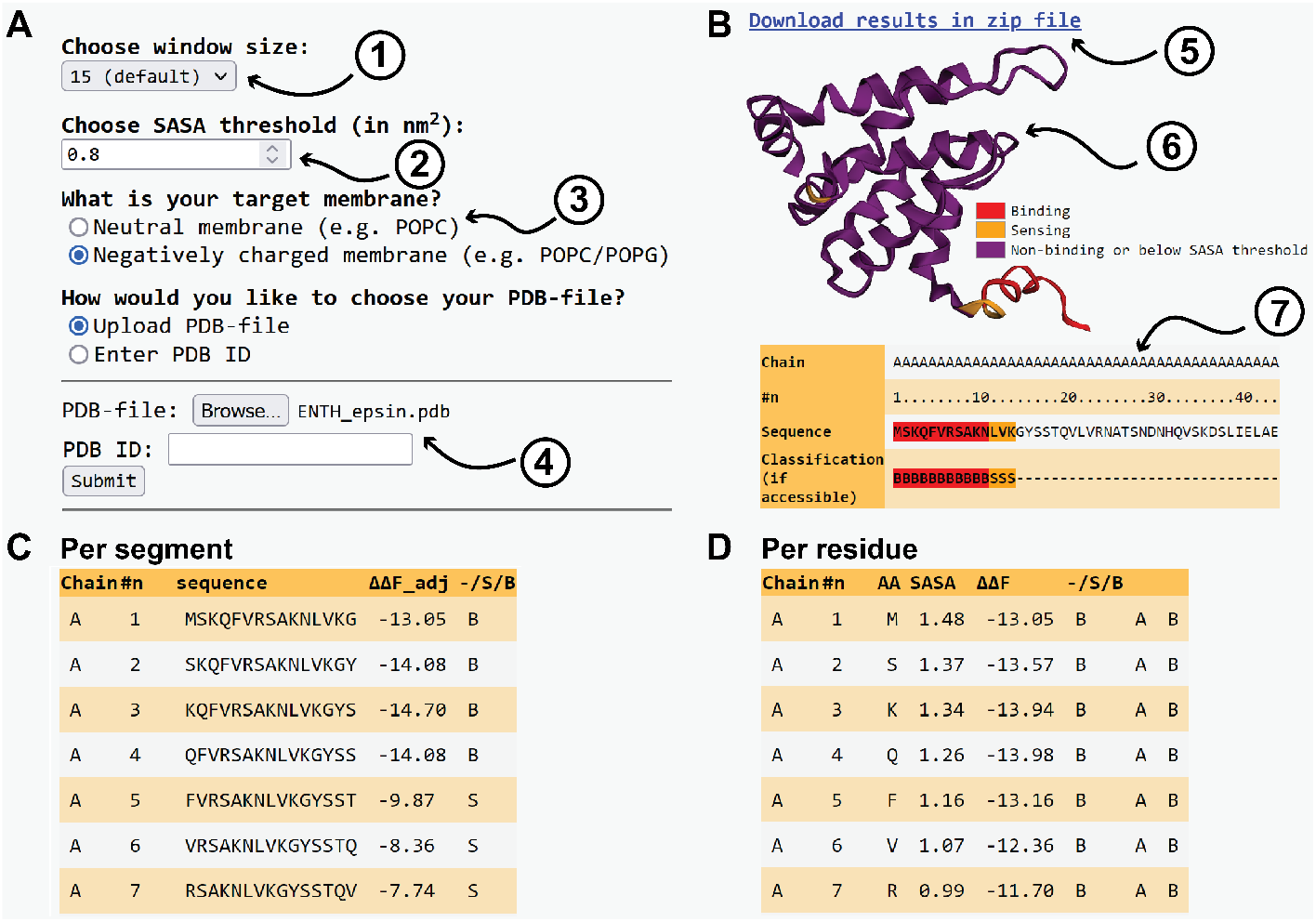
Example output for PMIpred’s protein module. **A)** The module takes four inputs: (1) The size of the sliding window (from 7-24 residues, default=15). (2) The SASA threshold (default=0.8 nm^2^), above which a residue is considered sufficiently solvent-accessible to contribute to membrane interactions. (3) The target membrane. (4) An uploaded PDB-file or PDB ID. **B)** Output includes: (5) A downloadable zip-file containing all output files that are also visualised on this page. (6) An interactive visualisation of the protein structure using 3Dmol (Rego and Koes [2015]). Predicted non-binding, sensing, and binding regions are coloured purple, orange, and red, respectively. (7) An output table containing the full protein sequence and the annotated classification: B=binder, S= sensor, -=non-binder or not accessible (SASA below threshold). **C)** A table containing the free energies and classifications of the individual segments that were screened. **D)** A more detailed look into the data underlying the table in B, including the SASA, ΔΔ*F* values, and classification per residue.

After submission, the results are displayed in the browser (Fig. 4B) and available to be downloaded as a zip-file. The processed protein is displayed interactively on the results page, using the aforementioned colouring: purple, orange, and red, for the non-binding (or buried) regions, predicted sensing domains, and membrane-binding domains, respectively. The data underlying this colouring are listed per segment (Fig. 4C) and per residue (Fig. 4D).

### 3.3 Cautionary remarks & recommendations

Before concluding this paper, we wish to point out several cases in which PMIpred’s results should be interpreted with some extra caution.

1. We note that shorter peptides have a higher tendency to be classified as binders (Fig. S5A). This effect can not be explained by poorer accuracy in the transformer’s prediction for shorter sequences, because the normalised error is constant over the full length range (7-24 residues, Fig. S5B). Alternatively, we suggest that this bias is likely introduced by our length extrapolation step (eq. 1) that intrinsically has a more pronounced effect for shorter peptides. Therefore, we advise users to interpret predictions on short peptides (≤ 12 residues) with some caution.
2. Our charge-correction (eq. 2) is based on a set of benchmark peptides from the experimental literature with charges ranging from *z* = −1 to *z* = +4 (van Hilten et al. [2023]). Predictions for peptides outside this charge range should be considered with some extra caution.
3. The interplay between membrane binding and peptide folding is also linked to the lipid composition of the membrane. This aspect is simplified in the MD simulations on which our model is trained, because we only used pure POPC and POPC/POPE membranes. If a protein’s membrane binding behaviour is linked to interactions with a specific lipid species, PMIpred may underestimate the membrane affinity.
4. The N- and C-terminal residues of proteins are more likely to be identified as (false) positives by the PMIpred protein screening module, for two reasons: (1) the termini of proteins tend to have a high solvent-accessibility. And, (2) the per-residue ΔΔ*F* is averaged over fewer positions, and thus possibly less accurate. Please note that the same applies to gaps in protein chains.
5. As mentioned in section 3.1, PMIpred always assumes an *α*-helical fold. Although our MD simulations suggest that folding (helical *vs* random coil) is a less determinant factor than AA composition (Fig. S4), we recommend users to take extra caution when working with that feature a known functional non-helical secondary structure. Of course, this also applies to non-helical domains in full-length proteins.
6. When using predicted 3D structures (e.g. by AlphaFold), we advise to only submit structures with high confidence – pLDDT ≥ 70 –, unless a section is known to be unstructured.
7. Protein-protein interaction sites may be falsely identified as membrane-binding regions by PMIpred, within the protein context. This misinterpretation is due to the fact that both phenomena are driven mostly by hydrophobic residues being exposed at the surface of the protein. For example, we observed this for the RAS-binding domain of PI(4,5)P_2_ 3-kinase, which included two exposed leucines L233 and L234 (see Table S3). We should emphasize here, however, that our method is based on peptide-data, and that the membrane-interaction predictions on the *peptide fragments* that make up a protein-protein interaction domain may still be valid.
8. Applying PMIpred to lipid-anchored proteins (e.g. KRAS in section S6) requires some extra caution, because our web server only considers the protein’s amino acids and neglects lipidic membrane anchors (e.g. Cys-linked farnesyl or palmitoyl moieties).
9. Although PMIpred is not designed for it, it can also be applied to transmembrane proteins, since the physicochemical properties driving membrane-surface binding and membrane internalisation are similar (hydrophobicity, mostly). We do note, however, that it may miss transmembrane regions for multi-pass proteins (e.g. GPCRs) due to the crowded protein environment and consequent low ‘solvent’-accessibility. Generally, we found that PMIpred behaves similar to DREAMM in this regard (Fig. S7). When working on transmembrane proteins, we recommend using PMIpred in conjunction with other web servers that are specifically developed for this (e.g. DeepTMHMM (Hallgren et al. [2022]), TMAlphaFold (Dobson et al. [2023]), or others).

## 4 Conclusions

We developed a transformer NN that is able to predict the relative binding free energy ΔΔ*F* for peptides interacting with (curved) membranes, only requiring an amino acid sequence as input. We implemented this model into a user-friendly web server, named PMIpred (https://pmipred.fkt.physik.tu-dortmund.de), where researchers can readily predict sensing free energies for any peptide, without requiring any simulation or coding experience.

As a second module in PMIpred, users can screen protein structures for putative curvature-sensing or membrane-binding motifs. Hereto, a sliding window is applied to the protein sequence to calculate a free-energy contribution for every residue. By coupling this to the solvent-accessible surface area (SASA) of that residue, the module is able to map and visualise the predicted membrane-interaction activity onto the protein structure. We applied this method to a comprehensive and diverse test set of known PMPs, and show that PMIpred is on par with the state-of-the-art tools DREAMM, PPM3, and MODA. In contrast to these other web servers, PMIpred provides *quantitative* predictions in terms of membrane-interaction free energies that can be directly interpreted biophysically, and additionally enables users to distinguish curvature-sensing from membrane-binding motifs. PMIpred also quantifies the contribution/weight of each individual amino acids to the predicted membrane binding, which guides the design of point mutations to experimentally probe protein-membrane interactions.

Since this method is trained on physics-based data (where experimental findings contribute only indirectly through the parametrisation of the MD force field), we argue that it should be able to characterise membrane-interaction behaviour from a broadly applicable fundamental perspective. In contrast, most current approaches are based on physicochemical descriptors, sequence ‘grammar rules’ that define known curvature-sensing motifs (e.g. ALPS (Drin et al. [2007])) but are unable to generalise their predictions to unrelated membrane-interacting motifs (e.g. *α*-synuclein and helix-0 of N-BAR and ENTH). When it comes to predicting the membrane interaction sites in PMPs, current methods are mainly data-driven, and thus rely on experimental findings on natural proteins. Although these tools are already remarkably powerful, we argue that the unique physics-informed predictions by PMIpred can additionally boost the performance of existing ensemble classifiers such as DREAMM, especially because it provides insights from a completely different angle and the data sources are orthogonal. Ultimately, by making PMIpred easily accessible for both computational and experimental biologists, we aim to accelerate the discovery and characterisation of curvature-sensing and membrane-binding peptides and protein motifs.

## Acknowledgments

The authors would like to thank Agata Witkowska for fruitful discussions and feedback. Art Hoti is thanked for thoroughly testing the web server.

## Author contributions

NvH and HJR designed the research and wrote the manuscript. NvH, NV, JM, and AB wrote the code. NV and AB designed, trained, and optimised the transformer model. NV and NvH designed the web server. CN was responsible for hosting the web server, and provided technical assistance.

## Data availability

The data and scripts underlying this article are available in https://github.com/nvanhilten/PMIpred.

