## Supplementary materials for "PMIpred: A physics-informed web server for quantitative Protein-Membrane Interaction prediction"

### S1 Methods: the transformer model

**Training data.** The transformer model was trained on the same as the data described in ref. (van Hilten et al. [2023]), comprising 53,940 unique sequences (all 24 residues long). The MD-calculated $\Delta\Delta F$ distribution for these peptides is shown in Fig. S1. All training data is available at `https://github.com/nvanhilten/PMIpred`.

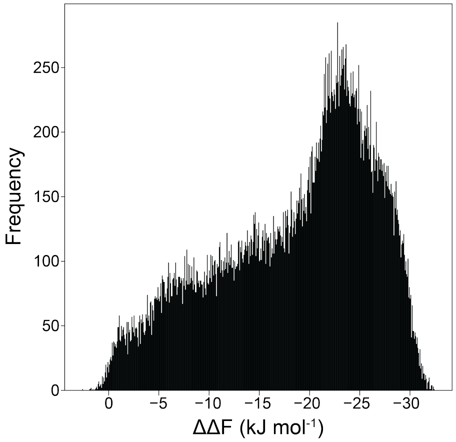


**Fig. S1** $\Delta\Delta F$ **distribution of the training data.** $\Delta\Delta F$ distribution of all 53,940 sequences in the training data.

**Data processing.** Within our NN, we use a tokeniser to assign a unique integer (1-20) to each AA type (Fig. S2A). Despite all training sequences being 24 AAs long already, we applied padding (adding a "0" token at the end) up to a fixed maximal length of 24 to enable the model to also process shorter sequences (as present in the test sets).

Because the position of an amino acid in a peptide sequence is relevant for its function, we added a vector containing this positional information to the token vector (Fig. S2A). Importantly, the weights linked to this so-called positional embedding, can be adjusted during training to tune the influence of the respective positions and AA types.

**Model architecture.** The final architecture of our transformer NN includes a series of 6 transformer blocks that use multi-head attention to learn patterns in the training data sequences (Fig. S2B). An output dense layer translates the encoded data into a float value; the predicted sensing free energy $\Delta\Delta F$.

**Model validation & testing.** Since the training set includes highly similar sequences from late iterations of the evolutionary algorithm, we validated the model using a more challenging target: 987 fully independent, randomly generated sequences (7-24 amino acids long), that were split into a validation set and a test set (494 and 493 sequences, respectively), keeping the latter completely separate from the training process. The $\Delta\Delta F$ values of these test

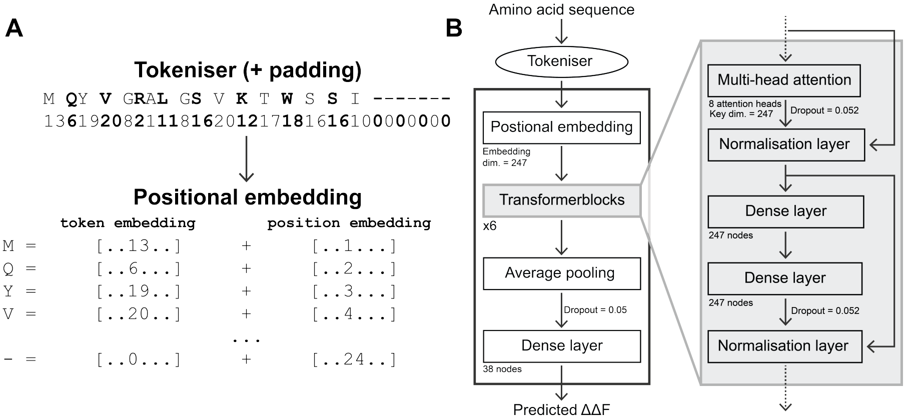


**Fig. S2 Transformer details. A)** Schematic description of how positional embedding is implemented in the transformer model. "[..X...]" represents a vector of a token or the position of that token in the sequence.

**B)** Architecture of the final transformer model.

set sequences, predicted by the final model, strongly correlated with the MD results ($R^2 = 0.80$, MSE= 1.58 kJ$^2$ mol$^{-2}$, RMSE= 1.26 kJ mol$^{-1}$, Fig. S3).

Additionally, to assess the performance of our model across the entire free-energy spectrum, we evaluated 96 sequences (24 residues each) that were not part of the training data and that ranged from 0.0 to -30.0 kJ mol$^{-1}$. Here, again, we found a close correlation between $\Delta\Delta F$ values predicted by the transformer model and MD-calculated free energies ($R^2 = 0.95$, MSE= 3.21 kJ$^2$ mol$^{-2}$, RMSE= 1.79 kJ mol$^{-1}$, Fig. 1B in the main text). For all test sets, the transformer model notably outperformed our previously trained CNN (van Hilten et al. [2023], Table S1). The validation set, test set, and full-range test set are all available at `https://github.com/nvanhilten/PMIpred`.

**Table S1**: **Model comparison.** $R^2$, mean squared error (MSE, in kJ$^2$ mol$^{-2}$), and root mean squared error (RMSE, in kJ mol$^{-1}$) between MD-calculated and NN-predicted $\Delta\Delta F$ for our previous CNN (van Hilten et al. [2023]) and our current transformer model on the three test sets (with $n$ sequences).

| | | | | CNN | | | Transformer | | |
|---|---|---|---|---|---|---|---|---|---|
| | **$\Delta\Delta F$ range** | **Sequence length** | **n** | **$R^2$** | **MSE** | **RMSE** | **$R^2$** | **MSE** | **RMSE** |
| **Validation set** | $0 \rightarrow -16$ | 7-24 | 494 | 0.657 | 2.915 | 1.707 | 0.769 | 1.829 | 1.352 |
| **Test set** | $0 \rightarrow -16$ | 7-24 | 493 | 0.663 | 2.924 | 1.710 | 0.800 | 1.578 | 1.256 |
| **Full range test set** | $0 \rightarrow -30$ | 24 | 96 | 0.941 | 6.225 | 2.495 | 0.954 | 3.211 | 1.792 |

**Hyperparameter optimisation.** During training, the transformer model was evaluated according to the mean squared error (MSE) between the prediction and the MD-calculated $\Delta\Delta F$ value for the 494 sequences in the independent validation set. The goal in training as to minimise this MSE.

The model was initialised using the glorot_uniform initialiser. ReLu activation was applied to every dense layer, except the final one. We used the Adamax algorithm as the optimisation method during training. Data shuffling

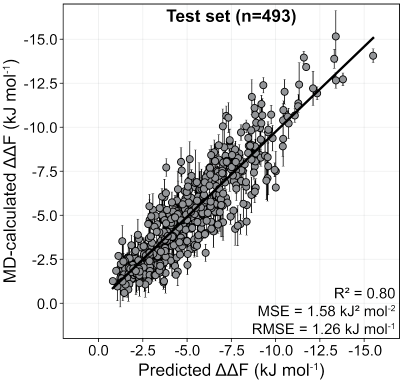


**Fig. S3 Correlation between transformer-predicted and MD-calculated $\Delta\Delta F$ values.** Correlation plot for 493 randomly generated peptide sequences (length 7-24, not part of the training data) in the test set. MD-calculated values are averages and standard deviations for 3 independent replicas (500 ns simulation per run, as described in ref. (van Hilten et al. [2023])).

was switched off. We applied a learning rate scheduler that started at $10^{-4}$ and linearly decreased the learning rate by $1.5 \cdot 10^{-6}$ per epoch until the 30$^{\text{th}}$ iteration, after which it was kept constant at $5.5 \cdot 10^{-5}$.

Hyperparameter testing was performed using Optuna's (Akiba et al. [2019]) TPESampler in combination with a HyperbandPruner that sampled hyperparameter settings from predefined ranges (see below), and pruned the training of poor performing models after 30 iterations (based on the validation MSE). The hyperparameters we considered were:

- The **embedding dimension**, i.e. the length of the vector in which the tokens and positions are embedded, was chosen between 64 and 256. This also determines the number of nodes in the dense layers within the transformer blocks.
- The number of **transformer blocks**, i.e. the 'depth' of the model, ranged between 4 and 12.
- The number of **attention heads** in the multi-head attention layer ranged between 6 and 12. Each head can recognise a pattern after training.
- The **dropout rate** within the transformer blocks (to prevent overtraining) ranged from 0.05 to 0.30.
- The **number of nodes in the final dense layer**, before generating output, was varied from 8 to 64.

Using fixed initialisation seeds and a batch size of 64, the best model (embedding dim.=247, 6 transformer blocks, 8 attention heads, dropout=0.052, and 38 nodes in the final dense layer, see Fig. S2B) had an MSE of 1.83 kJ$^2$ mol$^{-2}$ for the validation set and 1.58 kJ$^2$ mol$^{-2}$ for the test set (Fig. S3).

### S2 The role of secondary structure

We assumed $\alpha$-helical secondary structure in the MD simulations for all sequences in our training set. This assumption is based on the fact that most natural peptides or protein motifs that are known to bind to the surface of membranes indeed fold into an $\alpha$-helix to optimally orient fatty amino acids toward the hydrophobic membrane core and polar residues toward the water phase. Many peptides (e.g. the ALPS motif) actually *only* fold into this amphipathic helical secondary structure when bound to curved membranes, whereas they remain unstructured in solution (Drin and Antonny [2010]). Atomistic MD simulations of the N-terminal helix of endophilin's N-BAR domain on curved and flat membranes even showed that binding and folding are cooperative processes in the sense that $\alpha$-helical folding is promoted by the presence of hydrophobic lipid packing defects and that, in turn, these defects are stabilised by the folded peptide (Cui et al. [2011]).

Beyond the notion that $\alpha$-helicity and (curvature-specific) membrane binding are two intertwined phenomena, a practical reason for us to enforce helical folding is that the Martini force field (Souza et al. [2021]) we used in our simulations does not model changes in secondary structure. Instead, secondary structure elements are explicitly imposed through proper dihedrals between the backbone beads (Monticelli et al. [2008]).

To address the question whether folding would affect the sensing free energy $\Delta\Delta F$, we performed additional MD simulations of sequences sampled across the full applicability domain ($\Delta\Delta F$ roughly ranging from 0 to -30) with the peptides either explicitly modelled as an $\alpha$-helix or as an unstructured coil. When plotting the resulting free-energy differences for the two distinct secondary structures (Fig. S4), we find that:

1. They strongly correlate ($R^2 = 0.86$), which suggests that the activity is dominated by the general amino acid content rather than the secondary structure (as already suggested for antimicrobial peptides (Wimley [2010])).
2. Peptides with a low $|\Delta\Delta F|$ are more potent when modelled as a coil (left side of the dashed line in Fig. S4). We argue that this is due to the unstructured peptides being sufficiently flexible to be able to reorient their hydrophobic and hydrophilic residues to optimally match the membrane surface's amphipathic character.
3. Peptides with a high $|\Delta\Delta F|$ are more potent when modelled as a helix (right side of the dashed line in Fig. S4). This can be explained by a pre-organisation effect: the fixed helices may penetrate the membrane slightly further (thus inducing more tension) because the entropic penalty to do so is lower than for the unstructured peptides.
4. Peptides with a high hydrophobic moment (brighter colours in Fig. S4) yield higher $|\Delta\Delta F|$ values in a helical fold than as a coil. This is in line with the notion that there is a synergy between amphipathicity and (curvature-specific) membrane binding, as evident from the many examples in natural proteins.

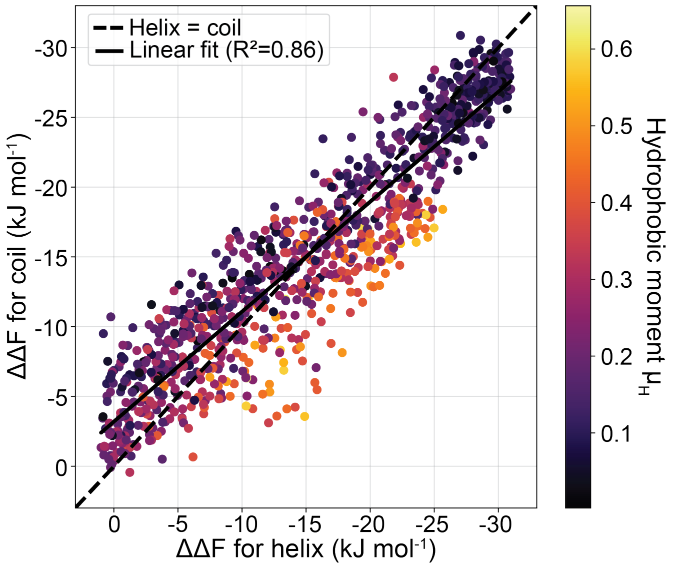


**Fig. S4 How secondary structure influences $\Delta\Delta F$.** Curvature-sensing free energies ($\Delta\Delta F$) for 992 peptide sequences that are modelled either as an $\alpha$-helix ($x$-axis) or as an unstructured coil ($y$-axis). Colors indicate the Eisenberg hydrophobic moment (Eisenberg et al. [1982], Fauchère and Pliška [1983]).

### S3 Protein test sets for benchmarking

**Table S2**: **The protein test set.** Overview of the protein structures used for benchmarking PMIpred against DREAMM, PPM3, and MODA (Fig. 3D in the main text). The top 19 proteins are the same as used by Chatzigoulas & Cournia's earlier benchmark (Chatzigoulas and Cournia [2022a]). The last 8 proteins are lipid packing defect sensors. The 27 proteins taken together are referred to as the "full test set". We used AlphaFold (Jumper et al. [2021], Varadi et al. [2022]) strucutres only in cases where there was no (complete) PDB file available. For every protein we used the exact same structure file (either from the PDB or AlphaFold) for the analysis. From left to right, columns give: The protein name (PI=phosphatidylinositol, PC=phosphatidylcholine); The PDB or AlphaFold (AF) database ID; The experimentally determined membrane-interacting residues; The corresponding membrane-interacting regions in our segment-based binary classification approach; References to the original literature.

| | Protein | ID | Residue(s) | Region(s) | Ref. |
|---|---|---|---|---|---|
| Chatzigoulas & Cournia's test set | Retinoid isomerohydrolase | 3fsn | 196-202, 264-271 | 183-212, 258-272 | Kiser et al. [2012] |
| | Voltage sensor toxin VSTx1 | 1s6x | 5-7, 27-30 | 1-31 | Jung et al. [2005], Ozawa et al. [2015], Lau et al. [2016] |
| | Cytotoxin 2 | 1ffj | 6-10, 22-39, 47-49 | 1-60 | Dubovskii et al. [2001] |
| | Sphingomyelinase C | 2dd (chain A) | 284-285 | 277-291 | Ago et al. [2006] |
| | Glycolipid transfer protein | 3rzn | 142 | 139-153 | Neumann et al. [2008], Ohvo-Rekilä and Mattjus [2011] |
| | Cholesterol-regulated Start protein 4 | 1jss (chain A) | 124 | 114-128 | Iaea et al. [2015] |
| | Ceramide transfer protein, PH domain | 2rsg | 33-36 | 24-38 | Sugiki et al. [2012] |
| | PI transfer protein beta isoform | 2a1l | 202-203 | 197-211 | Phillips et al. [2006] |
| | Cholesterol oxidase | 1coy | 81 | 79-93 | Chen et al. [2000] |
| | Cytochrome P450 3A4 | 1tqn | 44-47, 218-238 | 43-57, 208-237 | Baylon et al. [2013] |
| | 9-cis-epoxycarotenoid dioxygenase 1 | 3npe | 85-109, 222-237 | 83-112, 218-247 | Tan et al. [2001], Sui et al. [2013] |
| | Monoglyceride lipase MGLL | 3jw8 (chain A) | 179-186 | 166-195 | Karageorgos et al. [2017] |
| | Dihydroorotate dehydrogenase | 3w7r | 31-68 | 30-74 | Rawls et al. [2000], Costeira-Paulo et al. [2018] |
| | Phosphatase PTEN | 5bzz (chain A) | 260-269, 327-335 | 254-268, 325-339 | Lee et al. [1999], Nguyen et al. [2014] |
| | (S)-mandelate dehydrogenase | 6bfg | 177-215 | 168-227 | Xu and Mitra [1999] |
| | L-amino acid deaminase | 5hxw (chain A) | 345-355 | 339-368 | Ju et al. [2017] |
| | Intestinal fatty acid binding protein | 3akm (chain A) | 27 | 16-30 | Córsico et al. [2004], Falomir-Lockhart et al. [2006] |
| | PI(4,5)$P_2$ 3-kinase | 4ovu | 345-353, 378-379, 409-419, 720-729, 863-872 | 334-363, 378-393, 416-430, 716-730, 851-880 | Tubiana et al. [2022], Gabelli et al. [2010] |
| | PC transfer protein | 1ln1 | 184-193 | 173-202 | Feng et al. [2000] |
| Lipid packing defect sensor test set | ArfGAP1 | AF_Q8N6T3 | 197-231, 272-295 | 196-240, 271-300 | Bigay et al. [2005], Mesmin et al. [2007] |
| | N-BAR Bin1 | AF_Q9BTH3 (1-239) | 1-33 | 1-30 | Löw et al. [2008], Fernandes et al. [2008] |
| | ENTH epsin1 | 5onf | 4-15 | 1-15 | Ford et al. [2002], Stahelin et al. [2003], Kweon et al. [2006] |
| | Lipoprotein Lipase | AF_P11151 (35-478) | 416-426 | 410-439 | Lookene et al. [1997], Papadopoulou et al. [2023] |
| | Osh4/Kes1 | AF_A0A816BN28 | 1-23 | 1-30 | Drin et al. [2007] |
| | Barkor | AF_Q6ZNE5 | 468-490 | 466-492 | Fan et al. [2011] |
| | Pah1p (N-terminal domain) | AF_C7GU57 | 1-19 | 1-30 | Karanasios et al. [2010] |
| | $\alpha$-synuclein | 1xq8 | 2-23, 21-44 | 1-45 | Jensen et al. [2011], Pranke et al. [2011] |

### S4 Benchmarking results

**Table S3**: **Benchmarking results.** Segment-based binary classification results for three state-of-the-art methods (DREAMM (Chatzigoulas and Cournia [2022a,b]), PPM3(Lomize et al. [2022]), MODA (Kufareva et al. [2014])) and our PMIpred web server on the proteins from Table S2. TP = true positive, FP = false positive, TN = true negative, FN = false negative. The bottom three rows give the summed values for the respective test sets. The raw data is available at `https://github.com/nvanhilten/PMIpred`.

| | | DREAMM | | | | PPM3 | | | | MODA | | | | PMIpred | | | |
|---|---|---|---|---|---|---|---|---|---|---|---|---|---|---|---|---|---|
| | **Protein** | TP | FP | TN | FN | TP | FP | TN | FN | TP | FP | TN | FN | TP | FP | TN | FN |
| **Chatzigoulas & Cournia's test set** | Retinoid isomerohydrolase | 3 | 0 | 33 | 0 | 3 | 1 | 32 | 0 | 3 | 5 | 28 | 0 | 1 | 1 | 32 | 2 |
| | Voltage sensor toxin VSTx1 | 2 | 1 | 0 | 0 | 2 | 0 | 1 | 0 | 2 | 1 | 0 | 0 | 1 | 1 | 0 | 1 |
| | Cytotoxin 2 | 4 | 0 | 0 | 0 | 4 | 0 | 0 | 0 | 4 | 0 | 0 | 0 | 4 | 0 | 0 | 0 |
| | Sphingomyelinase C | 1 | 0 | 19 | 0 | 1 | 1 | 18 | 0 | 1 | 4 | 15 | 0 | 1 | 1 | 18 | 0 |
| | Glycolipid transfer protein | 1 | 1 | 12 | 0 | 1 | 1 | 12 | 0 | 1 | 1 | 12 | 0 | 1 | 3 | 10 | 0 |
| | Cholesterol-regulated Start protein 4 | 1 | 2 | 11 | 0 | 1 | 1 | 12 | 0 | 1 | 1 | 12 | 0 | 0 | 4 | 9 | 1 |
| | Ceramide transfer protein, PH domain | 1 | 2 | 4 | 0 | 1 | 2 | 4 | 0 | 1 | 3 | 3 | 0 | 1 | 1 | 5 | 0 |
| | PI transfer protein beta isoform | 1 | 1 | 16 | 0 | 1 | 1 | 16 | 0 | 1 | 1 | 16 | 0 | 1 | 3 | 14 | 0 |
| | Cholesterol oxidase | 0 | 2 | 31 | 1 | 1 | 3 | 30 | 0 | 0 | 3 | 30 | 1 | 0 | 0 | 33 | 1 |
| | Cytochrome P450 3A4 | 3 | 0 | 29 | 0 | 3 | 1 | 28 | 0 | 3 | 1 | 28 | 0 | 1 | 2 | 27 | 2 |
| | 9-cis-epoxycarotenoid dioxygenase 1 | 1 | 2 | 29 | 3 | 4 | 1 | 30 | 0 | 4 | 5 | 26 | 0 | 2 | 2 | 29 | 2 |
| | Monoglyceride lipase MGLL | 2 | 0 | 18 | 0 | 2 | 0 | 18 | 0 | 2 | 0 | 18 | 0 | 2 | 3 | 15 | 0 |
| | Dihydroorotate dehydrogenase | 2 | 1 | 21 | 1 | 3 | 0 | 22 | 0 | 3 | 1 | 21 | 0 | 1 | 1 | 21 | 2 |
| | Phosphatase PTEN | 1 | 1 | 18 | 1 | 0 | 3 | 16 | 2 | 1 | 4 | 15 | 1 | 2 | 5 | 14 | 0 |
| | (S)-mandelate dehydrogenase | 3 | 2 | 19 | 1 | 3 | 0 | 21 | 1 | 3 | 3 | 18 | 1 | 3 | 1 | 23 | 1 |
| | L-amino acid deaminase | 1 | 1 | 26 | 1 | 2 | 2 | 25 | 0 | 2 | 2 | 25 | 0 | 1 | 3 | 24 | 1 |
| | Intestinal fatty acid binding protein | 1 | 0 | 8 | 0 | 1 | 1 | 7 | 0 | 1 | 0 | 8 | 0 | 1 | 0 | 8 | 0 |
| | $PI(4,5)P_2$ 3-kinase | 1 | 5 | 75 | 6 | 0 | 1 | 79 | 7 | 3 | 5 | 75 | 4 | 3 | 6 | 74 | 4 |
| | PC transfer protein | 2 | 1 | 11 | 0 | 0 | 1 | 11 | 2 | 1 | 2 | 10 | 1 | 0 | 4 | 8 | 2 |
| **Lipid packing defect sensor test set** | ArfGAP1 | 4 | 12 | 6 | 1 | 1 | 2 | 16 | 4 | 5 | 13 | 5 | 0 | 5 | 3 | 15 | 0 |
| | N-BAR Bin1 | 1 | 1 | 13 | 1 | 0 | 5 | 9 | 2 | 2 | 7 | 7 | 0 | 2 | 0 | 14 | 0 |
| | ENTH epsin1 | 1 | 3 | 6 | 0 | 1 | 3 | 6 | 0 | 2 | 5 | 3 | 0 | 1 | 1 | 8 | 0 |
| | Lipoprotein Lipase | 2 | 1 | 22 | 0 | 2 | 5 | 18 | 0 | 2 | 11 | 12 | 0 | 2 | 4 | 19 | 0 |
| | Osh4/Kes1 | 1 | 2 | 25 | 1 | 1 | 4 | 23 | 1 | 1 | 5 | 22 | 1 | 1 | 4 | 23 | 1 |
| | Barkor | 2 | 10 | 21 | 0 | 0 | 6 | 25 | 2 | 2 | 22 | 14 | 0 | 2 | 6 | 25 | 0 |
| | Pah1p (N-terminal domain) | 2 | 5 | 0 | 0 | 2 | 1 | 4 | 0 | 2 | 3 | 2 | 0 | 2 | 3 | 2 | 0 |
| | $\alpha$-synuclein | 3 | 7 | 0 | 0 | 0 | 5 | 2 | 3 | 2 | 6 | 1 | 1 | 1 | 0 | 7 | 2 |
| | **Full test set** | 47 | 63 | 477 | 13 | 40 | 51 | 489 | 20 | 53 | 116 | 428 | 8 | 41 | 64 | 479 | 19 |
| | **Chatzigoulas & Cournia's test set** | 31 | 22 | 384 | 10 | 33 | 20 | 386 | 8 | 35 | 44 | 362 | 6 | 25 | 43 | 366 | 16 |
| | **Lipid packing defect sensor test set** | 16 | 41 | 93 | 3 | 7 | 31 | 103 | 12 | 18 | 72 | 66 | 2 | 16 | 21 | 113 | 3 |

The Matthews correlation coefficient (MCC, Matthews [1975], eq. S1) is a commonly used metric to score binary classifier predictions that is also suitable if the classes are differently sized (as is the case in our benchmark set, with more negatives than positives). It ranges from MCC = −1 (total disagreement between prediction and observation) to MCC = 1 (full agreement between prediction and observation, with MCC = 0 being the expected value for a random prediction.

$$\mathrm{MCC} = \frac{\mathrm{TP} \cdot \mathrm{TN} - \mathrm{FP} \cdot \mathrm{FN}}{\sqrt{(\mathrm{TP}+\mathrm{FP})(\mathrm{TP}+\mathrm{FN})(\mathrm{TN}+\mathrm{FP})(\mathrm{TN}+\mathrm{FN})}} \tag{S1}$$

The false discovery rate (FDR, eq. S2) is the fraction of positive observations (i.e. experimentally determined membrane-interacting regions) that are falsely predicted. The perfect classifier would have a FDR of 0%, i.e. none of the 'hits' are false positives.

$$\mathrm{FDR} = \frac{\mathrm{FP}}{\mathrm{FP}+\mathrm{TP}} \tag{S2}$$

The false omission rate (FOR, eq. S3) is the fraction of negative observations (i.e. regions that *do not* interact with membranes) that are falsely omitted. The perfect classifier would have a FOR of 0%, i.e. none of the experimentally determined membrane-interacting regions are missed.

$$\mathrm{FOR} = \frac{\mathrm{FN}}{\mathrm{FN} + \mathrm{TN}} \tag{S3}$$

The MCC, FDR, and FOR values that are reported in the main text are calculated from the summed values (bottom three rows in Table S3) using eq. S1, eq. S2, and eq. S3, respectively.

**Table S4**: **Scoring binary predictions.** MCC, FDR, and FOR values for three state-of-the-art methods (DREAMM (Chatzigoulas and Cournia [2022a,b]), PPM3 (Lomize et al. [2022]), MODA (Kufareva et al. [2014])) and our PMIpred web server on the proteins from Table S2. Values are calulcated using eq. S1-S3 on the summed values in the bottom three rows of Table S3. MCC data are also plotted in Fig. 3D in the main text.

| | | **DREAMM** | **PPM3** | **MODA** | **PMIpred** |
|---|---|---|---|---|---|
| **Full test set** | **MCC** | 0.49 | 0.46 | 0.44 | 0.44 |
| | **FDR** | 57% | 56% | 67% | 60% |
| | **FOR** | 3.5% | 4.7% | 2.3% | 4.4% |
| **Chatzigoulas & Cournia's test set** | **MCC** | 0.59 | 0.64 | 0.57 | 0.40 |
| | **FDR** | 42% | 38% | 53% | 61% |
| | **FOR** | 3.6% | 3.0% | 2.2% | 5.0% |
| **Lipid packing defect sensor test set** | **MCC** | 0.37 | 0.10 | 0.25 | 0.53 |
| | **FDR** | 72% | 82% | 80% | 57% |
| | **FOR** | 3.1% | 10.4% | 2.9% | 2.6% |

### S5 The effect on peptide sequence length on classification

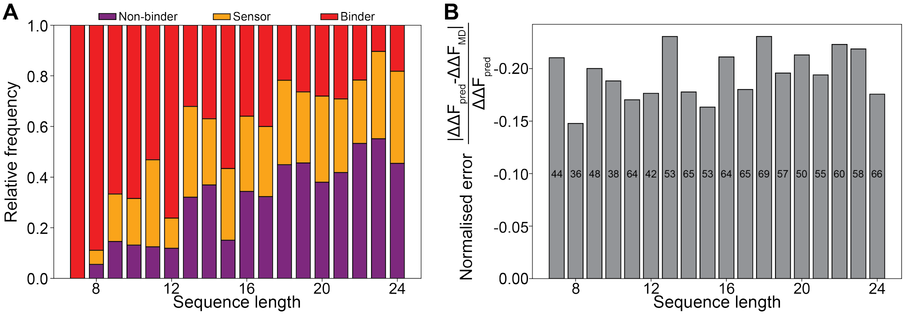


**Fig. S5 Sequence length affects classification. A)** The distribution of predicted non-binders, sensors, binders in the combined validation and test set (987 randomly generated sequences in total) per sequence length. Predictions were done for a negatively charged target membrane (i.e. using $\Delta\Delta F_{\mathrm{adj}}$). **B)** The normalised error for different sequence lengths in the combined validation and test set. Insets give the number of sequences per bin.

### S6 Lipid-anchored proteins

Lipid-anchored proteins are typically characterized by two features: a membrane-interacting protein domain and a covalently linked lipid anchor. For example, the GTPase KRAS features (1) a (polybasic) C-terminal region that interacts with anionic lipids in the membrane and (2) a Cys-linked farnesyl group that anchors it to the hydrophobic membrane interior.

We argue that the first of these features can be identified with PMIpred, whereas the second cannot. For KRAS in particular, we see that lysines K169 and K170 in the (variable) polybasic tail are correctly identified as membrane active when using a negatively charged target membrane in our model (Fig. X), even though it does not consider the farnesyl modification in this domain.

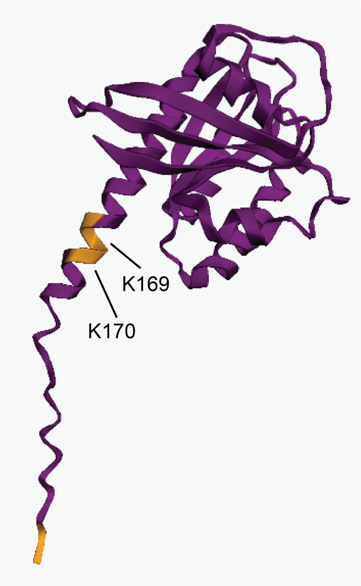


**Fig. S6 Predicting membrane-interaction sites in the lipid-anchored protein KRAS.** PMIpred result for KRAS. Orange indicates membrane-active (curvature-sensing) regions.

### S7 Transmembrane proteins

For monofunctional glycosyltransferase (a single-pass transmembrane (TM) protein), both PMIpred and DREAMM correctly predict most of the N-terminal helix as 'membrane binding' (Fig. S7A). In contrast, none of the TM helices from the $\beta_2$ adrenergic G protein-coupled receptor (GPCR) are identified as such by either web server (Fig. S7B), due to the crowded protein environment and consequent low 'solvent'-accessibility. The same becomes apparent when studying the homotetramer of the influenza A virus' M2 proton channel (Fig. S7C): both PMIpred and DREAMM prominently highlight the amphipathic helix (which is a known membrane binder and curvature inducer (Jung et al. [2019])), but not the TM part. In contrast, isolating one of the monomers renders the TM residues more 'solvent'-accessible, causing the full protein chain to be labeled membrane-interacting by both methods (Fig. S7D).

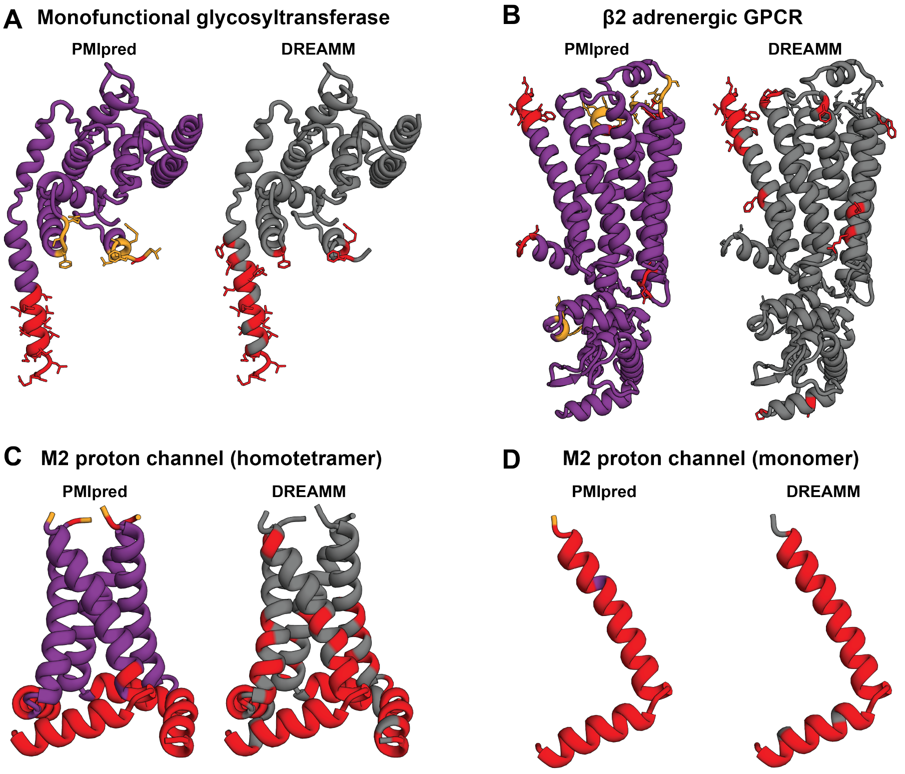


**Fig. S7 Predicting membrane-interaction motifs in TM proteins. A)** Predictions by PMIpred and DREAMM for monofunctional glycosyltransferase (PDB: 3vmt). Example taken from ref. (Chatzigoulas and Cournia [2022a]). **B)** Predictions by PMIpred and DREAMM for the $\beta_2$ adrenergic GPCR (PDB: 2rh1). **C)** Predictions by PMIpred and DREAMM for the influenza A virus' M2 proton channel (PDB: 2l0j) homotetramer and **(D)** monomer.
